# Using the drug-protein interactome to identify anti-ageing compounds for humans

**DOI:** 10.1101/438234

**Authors:** Matías Fuentealba Valenzuela, Handan Melike Dönertaş, Rhianna Williams, Johnathan Labbadia, Janet Thornton, Linda Partridge

## Abstract

Advancing age is the dominant risk factor for most of the major killer diseases in developed countries. Hence, ameliorating the effects of ageing may prevent multiple diseases simultaneously. Drugs licensed for human use against specific diseases have proved to be effective in extending lifespan and healthspan in animal models, suggesting that there is scope for drug repurposing in humans. New bioinformatic methods to identify and prioritise potential anti-ageing compounds for humans are therefore of interest. In this study, we first used drug-protein interaction information, to rank 1,147 drugs by their likelihood of targeting ageing-related gene products in humans. Among 19 statistically significant drugs, 6 have already been shown to have pro-longevity properties in animal models (p < 0.001). Using the targets of each drug, we established its association with ageing at multiple levels of biological actions including pathways, functions and protein interactions. Finally, combining all the data, we calculated a comprehensive ranked list of drugs that predicted tanespimycin, an inhibitor of HSP-90, as the top-ranked novel anti-ageing candidate. We experimentally validated the pro-longevity effect of tanespimycin through its HSP-90 target in *Caenorhabditis elegans*.

**Author Summary:** Human life expectancy is continuing to increase worldwide, as a result of successive improvements in living conditions and medical care. Although this trend is to be celebrated, advancing age is the major risk factor for multiple impairments and chronic diseases. As a result, the later years of life are often spent in poor health and lowered quality of life. However, these effects of ageing are not inevitable, because very long-lived people often suffer rather little ill-health at the end of their lives. Furthermore, laboratory experiments have shown that animals fed with specific drugs can live longer and with fewer age-related diseases than their untreated companions. We therefore need to identify drugs with anti-ageing properties for humans. We have therefore used computers to search for drugs that affect components and processes known to be important in human ageing. This approach worked, because it was able to re-discover several drugs known to increase lifespan in animal models, plus some new ones, including one that we tested experimentally and validated in this study. These drugs are now a high priority for animal testing and for exploring effects on human ageing.

## Introduction

Increasing life expectancy in developed countries is revealing advancing age as the primary risk factor for numerous diseases [1]. Thus, identifying interventions that can ameliorate the effects of ageing, and consequently delay, prevent or lessen the severity of age-related conditions, are needed. Extensive research in laboratory animals has demonstrated that the ageing process is malleable and that dietary, genetic and pharmacological interventions can improve health during ageing, extend lifespan and combat pathologies [2]. Furthermore, humans who lived to advanced ages show lower late-life morbidity (disease burden) than those who die earlier, indicating that compression of morbidity is achievable [3].

Although pharmacological interventions may prove to ameliorate the effects of ageing in humans, development of new drugs for this purpose would present significant difficulties, because of the need to treat healthy individuals in clinical trials over long periods for multiple outcomes. For this reason, it is more feasible to repurpose drugs already approved for specific diseases than to target ageing itself with new drugs [4,5]. With this goal in mind, researchers have begun to conduct human clinical trials to assess the anti-ageing properties of drugs approved to treat human medical conditions, and that extend lifespan and healthspan in animal models. Some examples include the antidiabetic drugs metformin (National Clinical Trial (NTC) number: NCT02432287) [6] and acarbose (NCT02953093), the immunosuppressant sirolimus (NCT02874924) and related compounds [7,8], and the natural compound resveratrol (NCT01842399). Two natural metabolites, the NAD precursors nicotinamide riboside (NCT02950441) and nicotinamide mononucleotide [9] are also being investigated. The development of computational methods to complement and accelerate this approach, by prioritising approved drugs that could ameliorate human ageing, is needed.

Several bioinformatic methods have been developed to identify potential geroprotective drugs. For instance, caloric restriction (CR) mimetics have been identified, by comparing genes differentially expressed in rat cells exposed to sera from CR rats and rhesus monkeys with gene expression changes caused by drugs in cancer cell lines [10]. Structural and sequence information on ageing-related proteins have been combined with experimental binding affinity and bioavailability data to rank chemicals by their likelihood of modulating ageing in the worm *Caenorhabditis elegans* and the fruit fly *Drosophila melanogaster* [11]. Drug-protein interaction information has also been used to predict novel pro-longevity drugs for *C. elegans*, by implementing a label propagation algorithm based on a set of effective and ineffective lifespan-extending compounds and a list of ageing-related genes [12]. A similar approach used a random forest algorithm and chemical descriptors of ageing-related compounds from the DrugAge database [13] together with gene ontology (GO) terms related to the drug targets [14]. Enrichment of drug targets has been assessed for a set of human orthologues of genes modulating longevity in animal models to identify new anti-ageing candidates [15].

Despite the increasing interest in drug-repurposing for human ageing, research has tended to focus on predicting life-extending drugs for animal models. However, the translation from non-mammalian species to humans is still a challenge, and certain aspects of ageing may be human-specific. Only a few studies have focused on data from humans. For instance, Aliper et al. (2016) [16] applied the GeroScope algorithm [17] to identify drugs mimicking the signalome of young human subjects based on differential expression of genes in signalling pathways involved in the ageing process. Another study by Dönertas et al. (2018) [18] correlated a set of genes up- and down-regulated with age in the human brain with drug-mediated gene expression changes in cell lines from the Connectivity Map [19].

In the present study, we rank-ordered drugs according to their probability of affecting ageing, by measuring whether they targeted more genes related with human ageing than expected by chance, by calculating the statistical significance of the overlap between the targets of each drug and a list of human ageing-related genes using a Fisher’s exact test [20]. Additionally, to enhance the power of the approach, we mapped the drugs’ gene targets and ageing-related genes to pathways (KEGG, Reactome), gene ontology terms (biological processes, cellular components, molecular functions) and protein-protein interactions, and repeated the analysis. We found that, independently of the data source used, the analysis resulted in a list of drugs significantly enriched for compounds previously shown to extend lifespan in laboratory animals. We integrated the results of 7 ranked lists of drugs, calculated using the different data sources, into a single list, and we experimentally validated the top compound, tanespimycin, an HSP-90 inhibitor, as a novel pro-longevity drug.

## Results

### Defining a dataset of drug-protein interactions and ageing-related genes

The drug-ageing association was inferred by comparing drug-gene interactions with gene-ageing associations. Fig 1 presents an overview of the procedure to prioritise the compounds. A dataset containing the interactions between drugs and proteins was built based on data from the STITCH database [21]. Only drugs targeting human proteins and successfully mapped to the DrugBank database [22] using the UniChem resource [23] were kept (Fig 1A). The dataset was composed of 18,393 interactions between 2,495 drugs and 2,991 proteins. More than half of the drugs (51.1%) in the dataset are approved for human use, 18.6% are in some phase of the approval process and 28.4% have been shown to bind to disease targets in experiments.

**Fig 1.**
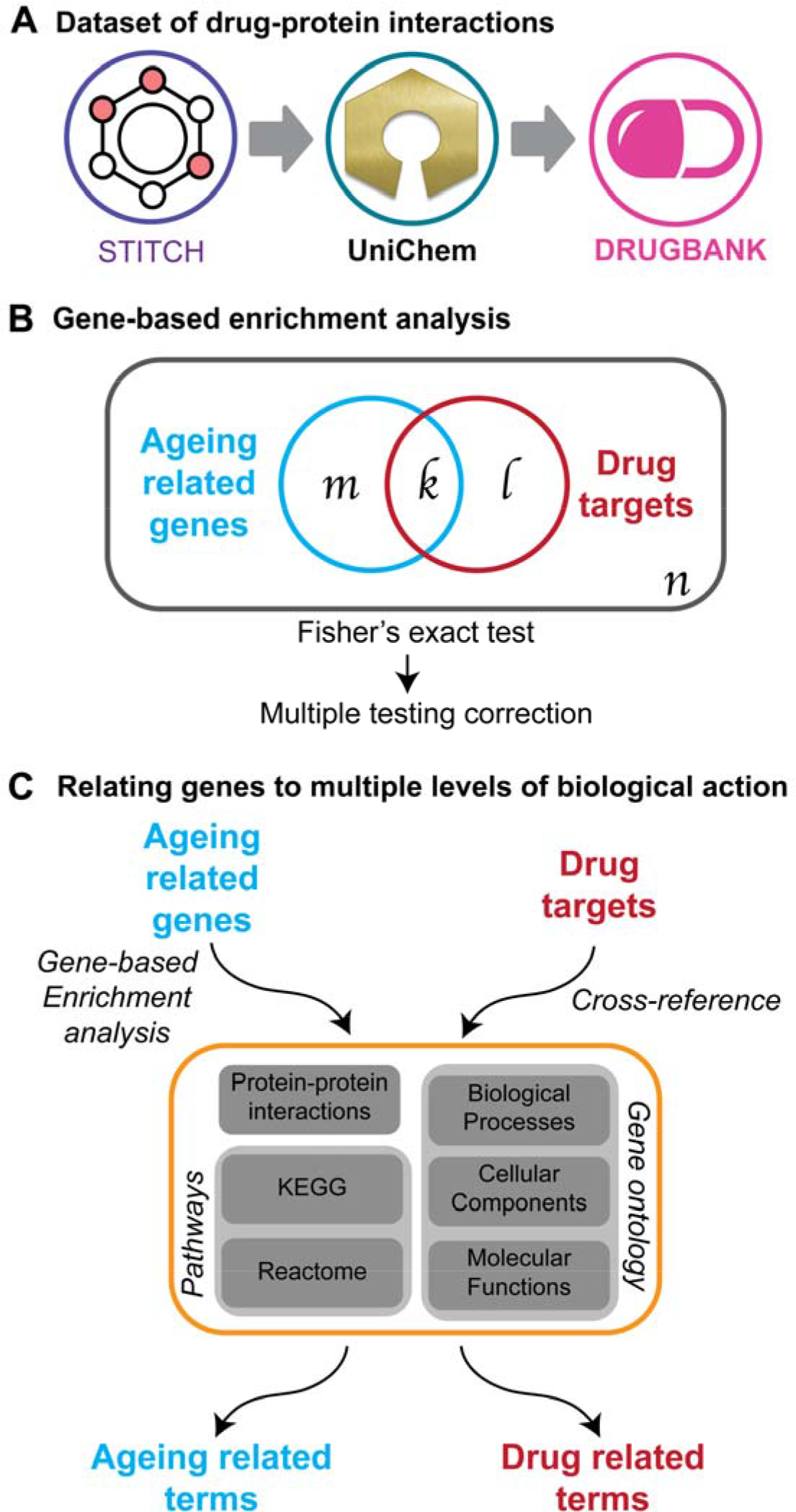
Overview of the methods used in this study to prioritise compounds likely to ameliorate ageing in humans. A) STITCH chemicals were mapped into DrugBank drugs using the UniChem resource programmatically. B) The significance of the drug-ageing inference was calculated using a Fisher’s exact test, which calculates the probability that the overlap between two samples (ageing-related genes and drug targets) drawn from the same universe is due to chance. This comparison was made at different biological tiers. C) Diagram of the procedure to expand the “gene” information into multiple biological levels. Ageing-related genes were mapped to other levels using an enrichment analysis, while the drugs’ targets were cross-referenced with the list of genes defining each annotation.

We obtained a set of ageing-related genes from the Aging Clusters resource [24]. A total of 1,216 ageing-related genes discovered in at least 2 among 4 categories of studies were selected. These 4 categories are human genes: i) changing expression with age or CR in different tissues ii) whose DNA methylation levels changes with age iii) associated with age-related diseases and iv) in manually curated databases of genes linked with longevity in genetic studies [25], associated with cellular senescence [26] or showing ageing-related effects in animal models in addition to evidence for a causative role in human ageing [27].

### Gene-based inference of drug-ageing associations

We determined if there was evidence supporting an association between drugs and ageing-related genes by calculating the statistical significance of the overlap between the gene targets of each drug and the ageing-related genes (Fig 1B). From the 1,147 drugs analysed, 19 were statistically enriched for ageing-related targets after multiple testing correction (Table 1, S1 tables). To assess the capability of the method to prioritise pro-longevity compounds, we compared the list of top-ranked compounds with the DrugAge database [13]. Six out of the 19 drugs have already been reported to significantly extend the lifespan of at least one model organisms (S2 text), while only 1 was expected by chance (p < 0.001). Additionally, using literature mining, we identified studies showing the association with longevity of cAMP analogues [28], selenium [29,30] and tanespimycin [31,32]. In contrast, we also found evidence for the DNA-mediated, pro-ageing (anti-longevity) effects of doxorubicin [33], cisplatin [34] and hydrogen peroxide [35]. We performed an interaction-based similarity analysis and found that the genotoxic compounds cluster separately from the other drugs, suggesting that they have a similar mechanism of action (S2 text). Similarities were also identified regarding the mechanisms of action of sorafenib and regorafenib, bexarotene and GW-501516, and sirolimus and ECGC, in agreement with previous studies [36].

**Table 1.** Drugs significantly enriched for ageing-related targets. The names of the drugs previously shown to extend lifespan in animal models are in bold and genotoxic molecules are in italic. The columns k(l) and m(n) are consistent with the diagram in Fig 1B. OR stands for odd-ratios and adj.p-value is the p-value adjusted for multiple testing.

| Drug name | Status | k(l) | m(n) | OR | p-value | adj.p-value |
| --- | --- | --- | --- | --- | --- | --- |
| **Resveratrol** | Approved | 66(150) | 388(2221) | 2.52 | 2.09E-08 | 1.82E-04 |
| Sunitinib | Approved | 18(12) | 436(2359) | 8.11 | 4.92E-08 | 2.15E-04 |
| **Genistein** | Investigational | 41(80) | 413(2291) | 2.84 | 6.40E-07 | 1.86E-03 |
| **Simvastatin** | Approved | 39(77) | 415(2294) | 2.80 | 1.53E-06 | 3.35E-03 |
| Tanespimycin | Investigational | 15(12) | 439(2359) | 6.71 | 2.64E-06 | 4.62E-03 |
| Regorafenib | Approved | 12(7) | 442(2364) | 9.16 | 4.43E-06 | 6.45E-03 |
| **Epigallocatechin gallate** | Investigational | 42(93) | 412(2278) | 2.50 | 5.96E-06 | 7.44E-03 |
| *Doxorubicin* | Approved | 34(67) | 420(2304) | 2.78 | 7.20E-06 | 7.87E-03 |
| Selenium | Approved | 14(12) | 440(2359) | 6.25 | 9.44E-06 | 9.17E-03 |
| **Celecoxib** | Approved | 23(36) | 431(2335) | 3.46 | 1.58E-05 | 1.38E-02 |
| Indole-3-carbinol | Investigational | 13(11) | 441(2360) | 6.32 | 1.83E-05 | 1.46E-02 |
| *Hydrogen peroxide* | Approved | 59(165) | 395(2206) | 2.00 | 2.85E-05 | 2.07E-02 |
| GW-501516 | Investigational | 9(5) | 445(2366) | 9.56 | 6.23E-05 | 3.82E-02 |
| Bexarotene | Approved | 10(7) | 444(2364) | 7.60 | 6.98E-05 | 3.82E-02 |
| Dorsomorphin | Experimental | 10(7) | 444(2364) | 7.60 | 6.98E-05 | 3.82E-02 |
| Sorafenib | Approved | 23(41) | 431(2330) | 3.03 | 7.25E-05 | 3.82E-02 |
| **Sirolimus** | Approved | 37(88) | 417(2283) | 2.30 | 7.42E-05 | 3.82E-02 |
| *Cisplatin* | Approved | 34(78) | 420(2293) | 2.38 | 8.39E-05 | 4.07E-02 |
| cAMP | Experimental | 36(86) | 418(2285) | 2.29 | 1.00E-04 | 4.60E-02 |

Although drugs interact directly with proteins, proteins do not act alone and interact with other proteins within pathways to perform different functions. Anti-ageing effects are likely to be mediated through altered pathway activity and function, and we therefore investigated if we could enhance the prediction of pro-longevity drugs using other biological annotations as comparators. Therefore, we calculated the pathways and gene functions enriched in ageing-related genes, together with the proteins that interact with them. A total of 82 KEGG and 54 Reactome pathways were enriched in this set of genes, as well as 1,177 biological processes, 69 cellular components and 103 molecular functions. In addition, we calculated that proteins interacted with the set of ageing-related genes. These terms, mapped at different biological levels, were defined as the set of ageing-related terms (Fig 1C – left). Equivalently, drugs were then associated with these terms through association with their targets using the list of genes defining each term according to the DAVID knowledgebase [37] and the biological database network [38]. This mapping procedure resulted in a set of terms from each data source related to each drug (drug-related terms) (Fig 1C – right).

### Drug-ageing association based on protein-protein interactions, gene ontology and pathways

Analogously to the gene-based association analysis, we calculated for each level if the overlap between ageing-related terms and drug-related terms was statistically significant using a Fisher’s exact test. This procedure generated 6 lists of ranked compounds in addition to the gene-based analysis (S1 tables). Notably, when we evaluated the correlation between the ranking of compounds in the different lists (Fig 2A), we observed a moderate correlation (Kendall’s coefficient of concordance W = 0.58, p-value = 1.02E-266). The highest correlations were observed between the results from biological processes and cellular components (Kendall’s tau = 0.51, p-value < 2.2E-16), while the lowest was observed between cellular components and genes (Kendall’s tau = 0.16, p-value = 3.289E-11).

**Fig 2.**
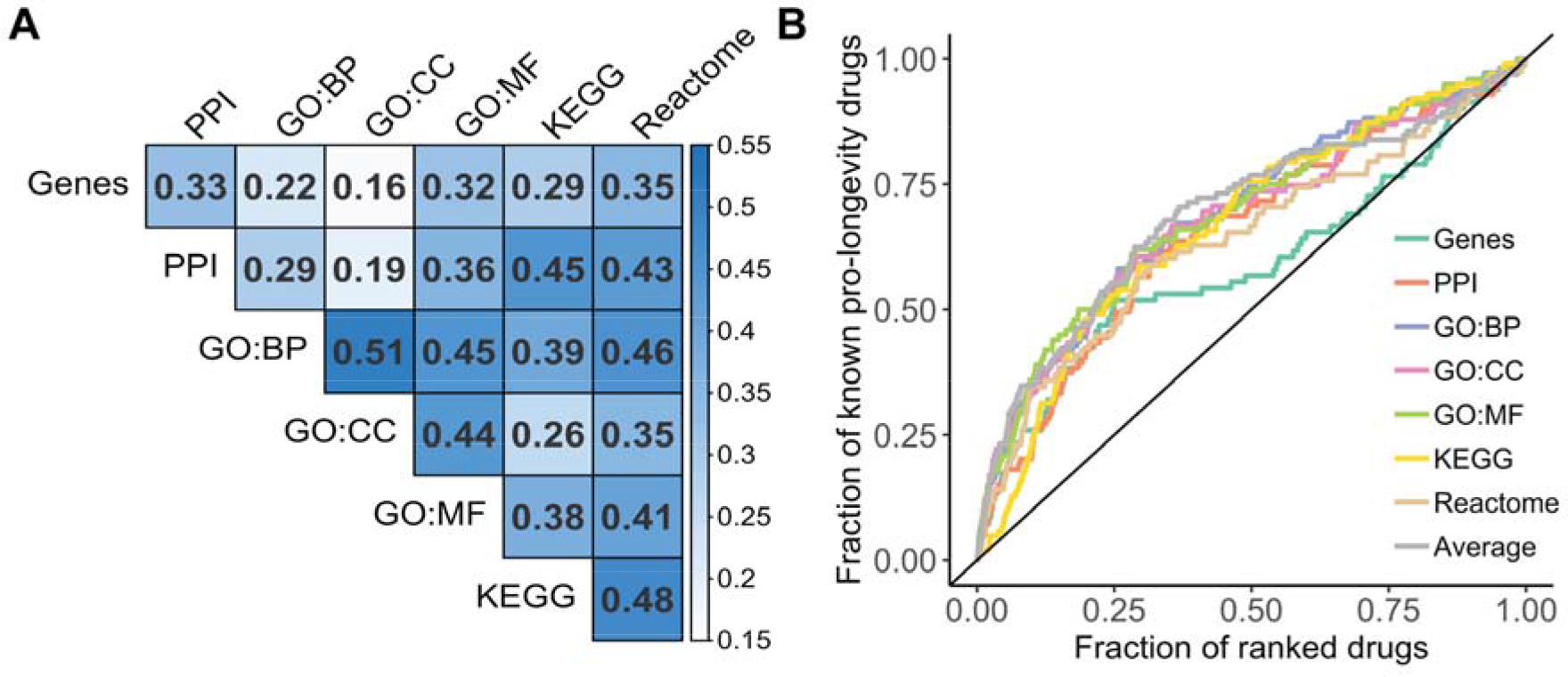
Comparison between the results using different data sources. A) Correlation between the ranked list of compounds. Boxes are coloured by the Kendall’s correlation coefficient. B) Enrichment curves for pro-longevity drugs. The results of each data source are displayed in lines with different colours. The enrichment expected by chance is shown as a diagonal line with AUC = 0.5.

Because in any enrichment analysis there is a potential for research bias, we performed random permutations to simulate the enrichment of each drug for a different set of terms on each level. None of the top-ranked drugs on each list ranked higher than in the analysis in more than 1.7% of the simulations (Table A in S2 text). We also quantified the capability of the strategy to prioritise pro-longevity compounds by calculating for each list the fraction of known pro-longevity compounds (ranked by p-value) among the fraction of drugs considered in each analysis (Fig 2B). The enrichment for pro-longevity compounds was quantified by calculating the area under the curve (AUC) generated by plotting these two variables. The maximum AUC was obtained when biological processes or molecular functions (AUC = 0.69) was used as the comparator (Table B in S2 text). The use of genes showed the lowest enrichment when non-statistically significant drugs are considered (AUC = 0.59), which suggests that the use of higher biological levels to calculate the inference improves the prediction capabilities, and that the use of genes leads to a loss power to rank drugs targeting a low proportion of ageing-related genes, which is observed in Fig 2B a loss of enrichment after 25% of the drugs were ranked. We evaluated if the AUCs were statistically significant by calculating the AUC from the simulations generated to quantify the research bias. The p-value for each curve was calculated by determining the number of simulated results with an AUC equal or higher than the analysis. All lists showed a higher enrichment than expected by chance (AUC > 0.5 and p-value < 0.05, Table B in S2 text). When we only considered the first 20 top-ranked drugs, we observed that using biological processes or cellular components to perform the comparison showed the highest proportion of pro-longevity drugs (45%), while only 2 pro-longevity drugs (10%) were found among the top 20 drugs when KEGG pathways are used.

Considering the lack of overlap between the ranked lists using the different data sources, we decided to integrate the results into a single list accounting for the complexity of multitiered effect of drugs by calculating their ranking average in the different analyses. The combination generated a list equally enriched as the maximum AUC obtained by the previous analysis (AUC = 0.69). Among the top 10 drugs with the best average ranking (Table 2, S1 tables), we found 3 drugs that have extended lifespan in animal models (trichostatin [39], geldanamycin [10] and celecoxib[40]). Half of these 10 drugs are classified as kinase inhibitors, while 8 are indicated as anti-cancer drugs and 7 are approved for human use.

**Table 2.** Top-ranked compounds using multiple levels of biological action. The names of the drugs previously shown to extend lifespan in animal models are in bold. The numeric values represent the ranking of the drugs when different sources of data (columns) are used. The last column is the ranking average (Avg.) for each drug in the 7 ranked lists.

| Drug name | Status | Genes | PPI | Gene ontology |  |  | Pathways |  | Avg. |
| --- | --- | --- | --- | --- | --- | --- | --- | --- | --- |
|  |  |  |  | BP | CC | MF | KEGG | Reactome |  |
| Tanespimycin | Investigational | 5 | 26 | 57 | 43 | 44 | 39 | 9 | 31.86 |
| Imatinib | Approved | 63 | 3 | 21 | 34 | 12 | 66 | 38 | 33.86 |
| Sunitinib | Approved | 2 | 1 | 59 | 31 | 31 | 56 | 63 | 34.71 |
| **Trichostatin** | Experimental | 83 | 41 | 19 | 54 | 13 | 41 | 52 | 43.29 |
| **Geldanamycin** | Investigational | 32 | 37 | 87 | 76 | 47 | 13 | 21 | 44.71 |
| Sorafenib | Approved | 16 | 68 | 11 | 15 | 8 | 155 | 42 | 45.00 |
| Dasatinib | Approved | 41 | 12 | 43 | 81 | 62 | 49 | 35 | 46.14 |
| Erlotinib | Approved | 27 | 6 | 93 | 85 | 71 | 64 | 7 | 50.43 |
| Etoposide | Approved | 23 | 11 | 20 | 90 | 32 | 120 | 67 | 51.86 |
| **Celecoxib** | Approved | 10 | 2 | 33 | 42 | 34 | 180 | 70 | 53.00 |

### The HSP-90 inhibitor tanespimycin as a novel pro-longevity drug

Leading the joint ranking was tanespimycin, also known as 17-AAG, a well-characterized HSP-90 inhibitor that has been shown to activate the transcription factor HSF-1 and induce a heat shock response in multiple model organisms [26]. As a proof-of-principle, we decided to investigate whether tanespimycin could activate HSF-1 and extend lifespan in the nematode worm *C. elegans*. To test the efficacy of tanespimycin dosing in *C. elegans*, we grew worms expressing mCherry under the control of an HSF-1 responsive promoter [41] on solid media plates containing various doses of tanespimycin. Worms were exposed to tanespimycin continuously from the first larval stage (L1) of development, or exclusively from the first day of adulthood. Worms grown continuously on tanespimycin plates exhibited a dose-dependent activation of the HSF-1 transcriptional reporter, starting at 25 µM and peaking at 100 µM (Fig 3A-B). Similarly, exposure to tanespimycin plates exclusively in adulthood resulted in significant activation of the HSF-1 reporter at 50 and 100 µM concentrations. No markers of toxicity were observed in any treatment groups, except for the 100 µM larval group, which were developmentally delayed by 24 hours and had a significantly reduced brood size (data not shown), consistent with chronic HSP-90 inhibition [42]. Together, these data demonstrate that tanespimycin activates HSF-1 in *C. elegans* and that treatment exclusively in adulthood is not associated with overt toxicity.

**Fig 3.**
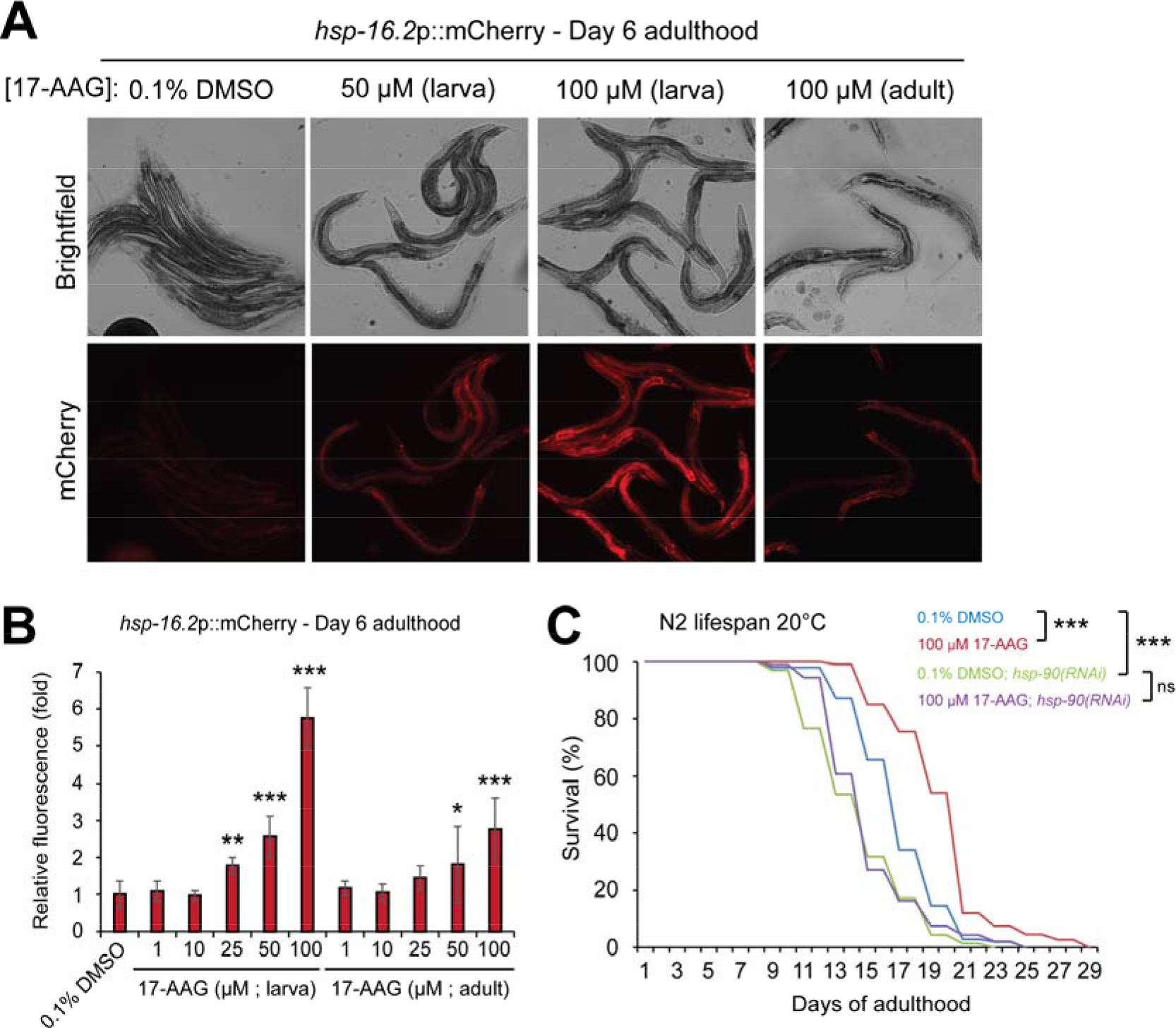
Pro-longevity effect of tanespimycin in *C. elegans*. A) Representative fluorescent images of day 6 adult, hsp-16.2p::mCherry transcriptional reporter worms, grown on plates containing 0.1% DMSO (vehicle) or different concentrations of tanespimycin (17-AAG) continuously from the first larval stage, or exclusively from the first day of adulthood onward. B) The relative fluorescent intensity of hsp-16.2p::mCherry worms grown on plates containing 0, 1, 10, 25, 50, or 100 µM tanespimycin (17-AAG) continuously from the first larval stage or exclusively from the first day of adulthood onward. Values plotted are the mean of at least 5 animals, and error bars represent the standard deviation from the mean. Statistical significance relative to the DMSO control group was calculated by ONE-WAY ANOVA with Tukey post analysis pairwise comparison of groups. * = p Ϗ 0.05, ** = p < 0.01, *** = p < 0.001. C) – Lifespan at 20 °C of N2 worms grown on plates containing 0.1% DMSO or 100 µM Tanespimycin (17-AAG) from the first day of adulthood onward in the presence or absence of hsp-90(RNAi). Statistical significance was calculated by Log-rank (Mantel-Cox) text. *** = p < 0.001. Treatment groups: 0.1% DMSO (n = 102, 14 censored, median lifespan = 17 days), 100 µM tanespimycin (n=107, 9 censored, median lifespan 21 days), 0.1% DMSO + hsp-90(RNAi) (n = 69, 30 censored, median lifespan = 15 days), 100 µM tanespimycin + hsp-90(RNAi) (n = 92, 22 censored, median lifespan = 15 days).

We next sought to determine whether tanespimycin treatment could extend lifespan in *C. elegans*. To circumvent potential longevity effects arising from delayed development and reproduction, we exposed worms to 100 µM tanespimycin plates from the first day of adulthood. Tanespimycin treatment significantly extended median and maximal lifespan compared to vehicle-treated controls (Fig 3C). To determine whether the effects of tanespimycin on lifespan require *hsp-90*, we also exposed worms to tanespimycin treatment in the presence of *hsp-90(RNAi)*. Consistent with previous reports, *hsp-90(RNAi)* significantly shortened *C. elegans* lifespan [43]. Furthermore, in the absence of HSP-90, tanespimycin treatment no longer increased lifespan compared to vehicle controls. These data suggest that tanespimycin treatment extends lifespan in an *hsp-90* dependent manner, but that severe depletion of HSP-90 is toxic to animals, despite the activation of protective stress responses.

## Discussion

This study was designed to infer and rank drugs matched to ageing at multiple levels of biological activity using a simple statistical test. In an initial gene-centric analysis, 19 drugs were identified as candidates expected to modulate ageing in humans. A major finding was that 6 of the statistically significant drugs, resveratrol, genistein, simvastatin, epigallocatechin gallate, celecoxib and sirolimus, have already shown lifespan-extending properties in experimental studies in model organisms. This statistically significant enrichment suggests that, despite its simplicity, the method is able to prioritise pro-longevity compounds. We then expanded the analysis to higher levels of biological complexity, and again found a statistically significant enrichment for pro-longevity drugs in all cases. The results of the analysis at different levels showed a moderate correlation. Compounds ranked high on average included trichostatin, geldanamycin and celecoxib, 3 drugs with pro-longevity effects in animal models [10,39,40]. The compound ranked highest on average was tanespimycin, an HSP-90 inhibitor, shown to acts as a senolytic agent by killing human senescent cells without affecting the viability of healthy cells [31] and to ameliorate disease phenotypes in *Drosophila* models of Huntington’s disease and spinocerebellar ataxia [32]. We found that tanespimycin treatment extended median (23%) and maximum (16%) lifespan in *C. elegans*, through its target HSP-90, possibly through the induction of cytoprotective pathways. Tanespimycin must act through more than one mechanism as a geroprotector, because cellular senescence has not been reported to occur in *C. elegans.*

Evidence from the literature supports the anti-ageing action of other drugs that we identified as potentially geroprotective. Dasatinib, a kinase inhibitor ranked 7^th^ on average, has been reported to induce apoptosis in senescent but not non-senescent primary human umbilical vein endothelial cells and preadipocytes [44]. Combination of dasatinib and quercetin, which also inhibits HSP-90, induced apoptosis specifically in senescent murine and human cells in vitro, improved cardiovascular function in aged mice, and decreased bone loss, neurological dysfunction and frailty in progeroid mice [45].

Three of the top 10 compounds from the combined ranked list have been previously proposed as anti-ageing candidates for humans using bioinformatic analysis. Specifically, tanespimycin, geldanamycin and trichostatin were among the 24 drugs predicted by Dönertas et al. (2018)[18] and Calvert et al. (2016)[10]. In contrast, we did not observe any overlap with the top results from Fernandes et al. (2016)[15] possibly due to the use of a different drug-protein interaction database (DGIdb [46]) or source of ageing data.

Similar enrichment-based methods that combine multiple levels of biological information have been used for drug-repurposing for Rheumatoid arthritis, Parkinson’s disease and Alzheimer’s disease [47,48], but not, to our knowledge, to identify anti-ageing drugs. Using annotated databases, our method evaluated the enrichment for pro-longevity of all compounds analysed, rather than only those with significant scores, and we observe that in all cases pro-longevity compounds are ranked higher than expected by chance. Although tanespimycin acts as a senolytic [31], and has been predicted to be geroprotective by two previous studies [10,18], we have demonstrated its effect on longevity experimentally.

A limitation of this study is that it is based on previous knowledge about drug-protein interactions, which for non-commonly studied drugs is incomplete. This may explain why we observed many anti-cancer and well-known drugs in our results. While we assessed this bias using permutations and we found no significant effect on our results, further research is needed to increase the drug-protein interactome data using, for example, high-throughput technologies like those currently available for kinases [49]. While we combined the results from the different data sources using a strategy based on ranks, we hypothesise that the integration of these results using other methods may lead to a list with a higher enrichment for pro-longevity drugs. Additionally, further experimental testing is required on the lists produced in this study, particularly those generated by using gene ontology terms, which presented the higher enrichment for pro-longevity drugs. An inherent limitation of inferred associations is that they do not provide information about the directionality of the effect, which in this case means that it is unknown if the drugs will deaccelerate ageing or the opposite. While we indirectly assessed this using an interaction-based similarity analysis between the drugs, resulting in clusters or pairs of drugs with similar mechanism of action, experiments should be conducted to determine the effects of each drug on ageing. Finally, a practical limitation is that we validated the results of this study using experiments in animal models although we used human data to perform the analysis. Although testing the effects of drugs on human ageing is challenging, progress is starting to be made. A clinical trial conducted by Mannick et al. (2018)[8] showed that pharmacological inhibition of the mammalian target of rapamycin in humans by dactolisib plus everolimus reduces the rate of infections in elderly people. Moreover, a recent short-term clinical trial of sirolimus established its safety in healthy individuals [50]. Similarly, supplementation of nicotinamide ribose, identified as a possible CR mimetic, stimulated NAD+ metabolism in healthy individuals aged 55 to 79 years [51]. Some mechanisms of ageing may be confined to humans and their near relatives, and ideally, the bioinformatic findings should be evaluated in humans, initially through genetic epidemiology and ultimately through clinical trials.

## Methods

### Data sources

#### Drug-protein interaction dataset

Chemical-protein interactions were extracted from the Search Tool for Interactions of Chemicals (STITCH) database 5.0 [21]. We chose this resource because it acts as a probabilistic network, by collecting interactions from multiple sources, including experiments, databases and a text-mining algorithm. Individual scores for each source are combined into an overall confidence score using a naive Bayesian formula defined as *Score* = 1 − ∏_*i*_(1 − *S*_*i*_), where S_i_ represents the confidence score for the source i. Later, because the Bayesian combination of scores can overestimate the effect of small individual contributions, the score is corrected for the probability of observing an interaction by chance. The overall confidence score ranges from 0 to 1, where a value of 0.4 or greater is considered as medium confidence, and a score equal to or higher than 0.7 is regarded as high confidence. To obtain a reliable set of interactions, we removed all interactions with a confidence score lower than 0.7. The database also maps the direction of each interaction, i.e. whether chemical acts on the protein or if the protein modifies the chemical (e.g. transformation of the chemical during a catalytic reaction). To confine the analysis to the actions of chemicals on proteins, only the cases where the chemical activates or inhibits a protein were retained. To focus on drugs in development or approved for human use, we filtered the chemicals in STITCH by the drugs in DrugBank 5.0 [22] using UniChem [23]. The InChi key for each drug was retrieved from PubChem (http://pubchemdocs.ncbi.nlm.nih.gov/pug-rest) and used to obtain the DrugBank identifiers via UniChem (https://www.ebi.ac.uk/unichem/info/webservices). The names of the drugs were obtained from the DrugBank vocabulary file, and the development status was acquired using the structure external links file. Finally, we mapped the Ensembl identifiers for each protein into HUGO Gene Nomenclature Committee (HGNC) approved gene names using Ensembl Biomart (version 91) [52]. All the chemicals included in this dataset will be referred as “drugs” or “compounds” throughout this article.

#### Drug-related terms

We mapped the targets of each drug in the drug-protein interaction dataset to multiple biological levels by using the information about the genes that define each level analysed. We downloaded the gene-centric definitions of *GO terms* and *Reactome pathways* from the DAVID knowledgebase [37]. Genes on each *KEGG pathways* were obtained using the biological database network (https://biodbnet-abcc.ncifcrf.gov/db/db2db.php)[38]. *Protein-protein interactions* were mapped directly using the STRING database [53]. Only proteins interacting with the set of ageing-related genes with a confidence equal of higher than 0.9 were considered.

#### Ageing-related genes

Genes present in manually-curated databases are more susceptible to research and reporting bias than those found in objective searches. Instead of selecting a set of ageing-related genes from a particular study or database, we used genes linked with ageing from the Ageing Clusters resource (https://gemex.eurac.edu/bioinf/age/). This repository contains the results of a network-based meta-analysis of human ageing genes [24] that considered 35 different datasets. The author classified the genes into the following 4 categories: curated ageing-related genes from databases such as GenAge [27], LongevityMap [25] and CSGene [26]; genes differentially expressed with age, regimes of CR or healthy ageing; age-related changes in the methylation of cytosine guanidine dinucleotides (CpGs) in the DNA; and genes associated with age-related diseases from databases such as the Human Gene Mutation [54] or the Human Phenotype Ontology [55]. To improve the reliability of the set of ageing-related genes and reduce research bias we considered only the genes present in at least two categories.

#### Ageing-related terms

Using the set of ageing-related genes we performed gene-based enrichment analysis to infer the function and pathways associated with ageing. *Gene Ontology (BP, CC, MF)* terms were calculated using the enrichGO function from the clusterProfiler package [56], using the Benjamini and Yekutieli [57] method for adjustment, a conservative correction that does not rely on the assumption that the test statistics are independent. P-value and q-value cutoff were set to 0.5 and for biological processes we consider the top 500 terms enriched. Enriched *KEGG pathways* were determined using the enrichKEGG function from the clusterProfiler package, using the same parameters used for the gene ontology enrichment. *Reactome pathways* were calculated using the function enrichPathway from the ReactomePA package [58]. *Protein-protein interactions*, were obtained using STRING [53] database.

### Statistical analysis to rank the drugs

Independently of the biological level, the drug-ageing associated was inferred by calculating the statistical significance of the drug-related terms and ageing-related terms using a Fisher’s exact test. Drugs were associated with ageing at the following biological levels: gene, pathways (KEGG, Reactome), functions (GO:BP, GO:CC, GO:MF) and indirect protein interactions. The universe was defined as all the terms on each level associated with at least one drug. Thus, drugs with a lower p-value modulate a higher proportion of ageing-related terms than that expected by chance. To control for the false discovery rate, we used the Benjamini and Yekutieli adjustment [57]. A p-value lower than 0.05 after multiple testing correction was considered significant.

### Measuring the impact of research bias

Some drugs have been more studied than others, which could bias the results towards drugs with a higher proportion of discovered targets. To evaluate the impact of this research bias, we randomly selected the same number of terms that were used as ageing-related terms 1,000 times, and we repeated the statistical analysis. Then we counted the times the statistically significant drugs appeared on the same or lower ranking. We expected that drugs associated with many terms would rank higher independently of the random set generated.

### Enrichment for pro-longevity drugs

Each of the drug lists generated were ranked by the p-values obtained from the statistical analysis. Then, we transformed the ranking of the drug into a value ranging from 0 to 1. A set of 142 pro-longevity drugs present in the DrugAge and DrugBank databases were used to determine the occurrence and ranking of pro-longevity compounds in the lists. The ranking was then scaled into a value between 0 to 1. The AUC between the variables describing the pro-longevity drugs and drugs analysed was calculated using the function AUC from the DescTools package (https://cran.r-project.org/package=DescTools). To measure its statistical significance, we calculated the AUC of the lists previously generated to measure the research bias, and we counted the number of simulations with an equal or higher AUC.

## Experimental procedure

### Worm husbandry and lifespan

N2 and TJ3002 (zSi3002*[hsp-16.2*p::*mCherry::unc-54*; *Cbr-unc-119(+)] II)*] hermaphrodite worms were maintained as previously described [59] at 20°C on 60 mm NGM plates. Plates were seeded with Escherichia coli (OP50) grown overnight in LB media. RNAi was essentially performed as previously described [60] with the slight modifications that bacterial cultures were induced with 5 mM IPTG for 3 hours following overnight growth in LB, and tetracycline was not included in plates or bacterial cultures.

### Tanespimycin dose-response test

Tanespimycin (Fisher Scientific) was solubilized in DMSO to stock concentrations of 1, 10, 25, 50, and 100 mM. 1 ml of DMSO or tanespimycin solutions were added to each litre of NGM media just prior to plate pouring to reach final concentrations of 1, 10, 25, 50, and 100 µM in plates. Plates were kept away from light, stored at 4°C, and used within 2 weeks of pouring. TJ3002 reporter worms were synchronised by bleaching and added to 0.1% DMSO or tanespimycin plates as L1s or as day 1 adults. Worms were transferred to fresh plates every day and then imaged on day 6 of adulthood using a Zeiss Apotome fluorescent microscope and Hamamatsu Orca Flash 4.0 camera. Brightness and contrast were adjusted linearly, and equally, for all images, using Adobe Photoshop CS6. Fluorescence intensity was measured under different conditions using ImageJ. Significance testing of differences in fluorescence intensity were calculated by ONE-WAY ANOVA with Tukey pair-wise comparison of groups using GraphPad Prism.

### Lifespan assays

Gravid N2 adults were bleached to release eggs, and L1 larvae were allowed to hatch overnight in M9 buffer without food. L1 worms were then added to plates seeded with bacteria expressing an RNAi control vector (L4440) and containing 0.1% DMSO. Worms were added to plates at a density of approximately 50 worms per plate. On the first day of adulthood (50h post plating L1s), worms were transferred to new 0.1% DMSO plates or 100 µM tanespimycin plates, seeded with L4440 or bacteria expressing dsRNA against hsp-90 (hsp-90(RNAi)). Worms were transferred to fresh plates every day during the first 7 days of adulthood and every other day thereafter. Worms were scored for survival every two days by gently prodding animals repeatedly with a platinum wire. Animals that failed to exhibit signs of movement or pharyngeal pumping were scored as dead. Animals that displayed internal hatching of progeny (“bagging”) or prolapse of intestine through the vulva (“rupturing”) were censored from our analysis. Median lifespans and significance testing between lifespans of different treatment groups were performed in GraphPad Prism using a Log-rank (Mantel-Cox) test.

## Acknowledgments

The authors thank the Partridge Lab and Dobril Ivanov for the support and helpful discussions. This work was funded by Comisión Nacional de Investigación Científica y Tecnológica - Government of Chile (CONICYT scholarship) (M.F.V.), BBSRC David Phillips Fellowship [BB/P005535/1] (J.L.), EMBL (H.M.D., J.M.T.), and the Wellcome Trust [WT098565/Z/12/Z] (L.P., J.M.T). H.M.D is a member of Darwin College, University of Cambridge.

